# Rab27 in tick extracellular vesicle biogenesis and infection

**DOI:** 10.1101/2023.11.02.565357

**Authors:** L. Rainer Butler, Nisha Singh, Liron Marnin, Luisa M. Valencia, Anya J. O’Neal, Francy E. Cabrera Paz, Dana K. Shaw, Adela S. Oliva Chavez, Joao H.F. Pedra

## Abstract

**Background:** The blacklegged tick, *Ixodes scapularis*, transmits most vector-borne diseases in the United States. It vectors seven pathogens of public health relevance, including the emerging human pathogen *Anaplasma phagocytophilum*. Nevertheless, it remains critically understudied when compared to other arthropod vectors. *I. scapularis* releases a variety of molecules that assist in the modulation of host responses. Recently, it was found that extracellular vesicles (EVs) carry several of these molecules and may impact microbial transmission to the mammalian host. EV biogenesis has been studied in mammalian systems and is relatively well understood, but the molecular players important for the formation and secretion of EVs in arthropods of public health relevance remain elusive. RabGTPases are among the major molecular players in mammalian EV biogenesis. They influence membrane identity and vesicle budding, uncoating, and motility.

**Methods:** Using BLAST, an *in-silico* pathway for EV biogenesis in ticks was re-constructed. We identified Rab27 for further study. EVs were collected from ISE6 tick cells after knocking down *rab27* to examine its role in tick EV biogenesis. *I. scapularis* nymphs were injected with small interfering RNAs to knock down *rab27* then fed on naïve and *A. phagocytophilum* infected mice to explore the importance of *rab27* in tick feeding and bacterial acquisition.

**Results:** Our BLAST analysis identified several of the proteins involved in EV biogenesis in ticks, including Rab27. We show that silencing *rab27* in *I. scapularis* impacts tick fitness. Additionally, ticks acquire less *A. phagocytophilum* after *rab27* silencing. Experiments in the tick ISE6 cell line show that silencing of *rab27* causes a distinct range profile of tick EVs, indicating that Rab27 is needed to regulate EV biogenesis.

**Conclusions:** Rab27 is needed for successful tick feeding and may be important for acquiring *A. phagocytophilum* during a blood meal. Additionally, silencing *rab27* in tick cells results in a shift of extracellular vesicle size. Overall, we have observed that Rab27 plays a key role in tick EV biogenesis and the tripartite interactions among the vector, the mammalian host, and a microbe it encounters.

## Background

The most medically relevant arthropod vector in the United States is *Ixodes scapularis*, the blacklegged or deer tick [1]. *I. scapularis* is responsible for spreading seven known human pathogens, encompassing bacteria, viruses, and parasites. In 2019, the Centers for Disease Control and Prevention (CDC) reported over 50,000 cases of tick-borne illness including 34,000 cases of Lyme disease and over 5,000 cases of Human Granulocytic Anaplasmosis [2]. Current estimates have calculated that Lyme disease alone costs the United States economy around $345 to $968 million per year [3]. Furthermore, the burden of vector-borne diseases on the healthcare system is expected to increase in the coming years [4, 5, 6, 7].

Ticks are hematophagous or blood feeding arthropods. When a tick takes a blood meal, it introduces its saliva into the wound along with any microbes it is carrying. Tick saliva has long been studied and is known to contain a variety of anesthetics, anti-coagulants, and anti-inflammatory compounds [8, 9, 10, 11]. However, the mechanism of delivery of these compounds was unknown until recently. It was found that ticks cells secrete extracellular vesicles (EVs) in their saliva [12]. Tick EVs have been shown in microbial transmission and to enable arthropod fitness [12, 13, 14]. Despite their role in arthropod fitness and disease, the molecular mechanisms and proteins involved in tick extracellular vesicle biogenesis remain poorly defined.

The bulk of EV biogenesis research has been completed in mammalian systems. In mammals, a variety of proteins participate in forming EVs including endosomal sorting complex required for transport (ESCRT) proteins, tetraspanins, soluble N-ethylmaleimide-sensitive factor attachment receptor (SNAREs), and Rab-GTPases [15]. These proteins aid in the transition from early to late endosome and the invagination of the endosome, which creates intraluminal vesicles (ILVs) and the multivesicular endosome (MVE) [16, 17, 18]. The MVE is then trafficked to and fuses with the plasma membrane, releasing EVs into the extracellular space. Once EVs are released, they are known to function in cell-to-cell communication, by transporting a variety of molecules. Additional evidence shows that components of the EV biogenesis pathway can be manipulated by intracellular bacteria and are important for transmission of arthropod-borne pathogens in other systems [12, 13, 19, 20, 21]. The function of EVs has not been thoroughly explored in arthropod biology. Although it is known that ticks and tick cells secrete EVs, it remains elusive how EV biogenesis occurs in ticks. Here, we indicate that silencing *rab27* in *I. scapularis* impacts arthropod fitness and microbial infection.

## Methods

### Mice and ticks

C57BL/6J mice were obtained from the University of Maryland Veterinary Resources. All mice used were between 6-8 weeks of age. *I. scapularis* nymphs were supplied by the Oklahoma State University and the University of Minnesota breeding colonies. Ticks were housed upon arrival at 23°C with 85% relative humidity and a 12/10-hour light/dark photoperiod cycle. All mouse experiments were performed according to the protocols approved by the Institutional Biosafety (IBC-00002247) and Animal Care and Use Committee (IACUC-01121014) at the University of Maryland School of Medicine and complied with National Institutes of Health (NIH) guidelines (Office of Laboratory Animal Welfare [OLAW] assurance number A3200-01).

### Tick cell culture

The *I. scapularis* ISE6 cell line was cultured in L15C300 medium supplemented with 10% heat-inactivated fetal bovine serum (FBS, 24 MilliporeSigma), 10% tryptose phosphate broth (BD), and 0.1% bovine lipoprotein concentrate (MP Biomedicals) at 34°C [22]. Tick cells were grown to confluence and sub-cultured in capped T25 flasks (Greiner bio-one). All cell cultures were verified by PCR to be *Mycoplasma*-free (Southern Biotech).

### A. phagocytophilum culture

*A. phagocytophilum* strain HZ was cultured in vented T25 flasks (CytoOne) containing HL-60 cells. HL-60 cells were cultured in 20 ml of RPMI medium, supplemented with 10% Fetal Bovine Serum and 1x Glutamax. *A. phagocytophilum* infected cells (500 μl) were added to 5 ml of uninfected cells at 1 to 5 × 10^6^ cells/ml diluted in 24.5 ml of media. The percentage of infection was monitored by the Richard-Allan Scientific™ three-step staining (Thermo Fisher Scientific). Infected cells were spun onto microscope slides with a Cytospin (Thermo Scientific). Cells were visualized by light microscopy with an Axioskop microscope (Zeiss). Bacteria were either used for experiments or sub-cultured once cultures had reached >90% infection. Bacterial numbers were estimated using the number of infected HL-60 cells × 5 morulae/cell × 19 bacteria/cell.

### Bioinformatics

Tick homologs of protein sequences involved in EV biogenesis in humans, mice, and *Drosophila melanogaster* were identified and compared using the NIH Basic Local Alignment Search Tool for proteins (BLASTp). Proteins with Expect (E)-values below 1×10^−5^ and greater than 80% query coverage were selected. The homologs identified were used to construct a putative EV biogenesis pathway for ticks.

### RNA interference

Small interfering RNAs (siRNAs) and scramble RNAs (scRNAs) were designed based on the sequence of tick *rab27* that was identified during the construction of the *in-silico* EV biogenesis pathway in ticks. siRNAs were designed using BLOCK-iT RNAi designer (https://rnaidesigner.thermofisher.com/rnaiexpress/), and scRNA was designed using InvivoGen siRNA Wizard Software (https://www.invivogen.com/sirnawizard/scrambled.php). Both siRNAs and scRNAs were blasted against the *I. scapularis* genome to confirm specificity and avoid off target effects. All siRNA primers can be found in Table S1.

For *in vitro* experiments, si-*rab27* and sc*-rab27* were synthesized by MilliporeSigma with dTdT overhangs. ISE6 cells were plated at 5×10^5^ cells per well (24 well plate) or 1×10^6^ cells per well (6 well plate). siRNAs (1 μg per ml) were nucleofected into ISE6 cells using the 4D-Nucleofector System (Lonza Bioscience). Tick cells were centrifuged at 100 x g for 10 minutes to pellet the cells. The pellet was washed with 10 ml dPBS, resuspended in SF buffer (Lonza Bioscience) and si*-rab27* or sc-*rab27* was added to the suspension. The nucleofection mix was added to a multiwell cuvette, inserted into the nucleofector, and pulsed using pulse condition EN150. The cells were then rested in the cuvette for 10 minutes post-nucleofection before being added to pre-warmed L15C300 complete media and seeded for experiments.

For *in vivo* experiments, si-*rab27* and sc*-rab27* were synthesized using the Silencer siRNA construction kit (Thermo Fisher Scientific) using the primers in Table S1. *I. scapularis* nymphs were microinjected with 20-40 ng of si*-rab27* or sc*-rab27*. Nymphs were rested for 24 hours before placement on mice.

### EV depleted medium

L15C300 medium was supplemented with 5% fetal bovine serum (MilliporeSigma), 5% tryptose phosphate broth (BD), and 0.1% lipoprotein concentrate (MP Biomedicals). Medium was cleared from EVs by ultracentrifugation at 100,000 ×g for 18 h at 4°C in a LE-80 ultracentrifuge (Beckman Coulter) with a 60Ti rotor [13]. The absence of EVs was confirmed by determining the particle size distribution with a NanoSight NS300 (Malvern Panalytical) for nanoparticle tracking analysis (NTA). If EVs were present, the medium was subjected to a second ultracentrifugation at 100,000 xg for 18 h at 4 °C. EV-free medium was sterilized by passing the content through a 0.22-μm Millipore Express® PLUS (MilliporeSigma).

### EV collection

ISE6 cells were nucleofected with si-*rab27* or sc-*rab27* as described above. After 72 hours, the L15C300 medium was replaced with EV depleted L15C300 medium. EV collection was carried out as described previously [13]. Briefly, the EV depleted medium was collected from cell cultures and cleared of any live cells by centrifugation at 300 × g for 10 min at 4 °C. Live cell pellets were resuspended in 1 mL of Trizol and saved for RNA extraction and qPCR. Dead cells were removed by a second centrifugation at 2000 × g for 10 min at 4°C. The supernatant was collected, and apoptotic bodies were removed by a third centrifugation at 10,000 × g for 30 min at 10 °C. To reduce the number of EVs >200 nm in size, the supernatant was filtered through a 0.22-μm Millipore syringe filter (Millipore-Sigma). EVs were pelleted by ultracentrifugation (100,000 × g) for 18 h at 4°C. Supernatant was discarded and EVs were resuspended in 1:500 in 1x PBS for NTA analysis.

### EV quantification

EV concentration and sizes were determined using a NanoSight NS300 (Malvern Panalytical) with NTA software version 3.0. The mean of the size generated in the NTA reports was used to calculate the average size of the EVs in each sample. Data was analyzed using GraphPad Version 10.0.3 from Prism.

### Quantitative reverse transcriptase polymerase chain reaction (qRT-PCR)

The PureLink RNA Mini kit (Invitrogen) was used to extract RNA from cells or ticks preserved in Trizol. cDNA was synthesized with the Verso cDNA Synthesis Kit (Thermo Fisher Scientific). qRT-PCR was performed with the CFX96 Touch Real Time PCR Detection System (Biorad). For bacterial acquisition experiments, *A. phagocytophilum 16S rRNA* gene expression was measured by absolute quantification and normalized to *I. scapularis actin*. Copy numbers for *A. phagocytophilum* and *I. scapularis* were calculated from a standard curve. Levels of tick *rab27* gene expression was measured by relative quantification and normalized to *I. scapularis actin*. The fold changes in gene expression were calculated using the 2−ΔΔC T method. Levels of gene expression were measured using Power SYBR® Green PCR master mix (Thermo Fisher Scientific). Amplifications were done using the following conditions: a denaturation cycle of 10 mins at 95 °C, followed by 40 cycles of denaturalization for 15 s at 95 °C, amplification at three different annealing temperatures (*rab27*: 58 °C; *Actin*: 57 °C; *A. phagocytophilum* 16S: 54°C) for 1 min. Primers were used at a final concentration of 400 ηM each and 2 μl of cDNA were used as template. The specificity of the products was determined by single peaks in the melting curves. Primers were designed using the PrimerQuest Tool (IDT). All primers used in this study can be found in Table S1.

### Tick fitness experiments

*I. scapularis* nymphs were microinjected with 20-40 ng of si*-rab27* or sc*-rab27*. Nymphs were rested for 24 hours. After 24 hours, the silenced or scrambled nymphs were placed on C57BL/6 J for 20 minutes and allowed to feed for 3 days. Ticks were recovered from a water trap (fully engorged) or were removed with forceps (partially engorged). The weight of the ticks was measured using a Pioneer™ analytical balance (OHAUS) and the differences in engorgement were evaluated. Ticks were stored at −80 °C until RNA extraction.

### *A. phagocytophilum* acquisition experiments

One week prior to nymph placement, 6–8-week-old male C57BL/6J mice were intraperitoneally (i.p.) injected with HL-60 cells containing *A. phagocytophilum* (1 × 10^7^ bacteria/injection) resuspended in 100 μL 1X PBS. At six days post infection (d.p.i.), *I. scapularis* nymphs were microinjected with 20-40 ng of si*-rab27* or sc*-rab27*. Nymphs were rested for 24 hours. At 7 d.p.i., the silenced or scrambled nymphs were placed on C57BL/6J for 20 minutes and allowed to feed for 3 days. Ticks were removed from mice with forceps, and fallen ticks were recovered from a water trap. Tick attachment was calculated based on contingency using Fisher’s exact test. The weight of the ticks was measured using a Pioneer™ analytical balance (OHAUS) and tick engorgement was evaluated. Ticks were stored at −80 °C until RNA extraction.

### Statistical analysis

Statistical significance of *rab27* silencing, vesicle concentration, and weight were assessed with an unpaired *t* test with Welch’s correction. Tick attachment was calculated based on contingency using a Fisher’s exact test. We used GraphPad PRISM® (GraphPad Software version 10.0.3) for all statistical analyses. Outliers were detected by a Graphpad Quickcalcs program (https://www.graphpad.com/quickcalcs/Grubbs1.cfm).

## Results

### Building an *in silico* extracellular biogenesis pathway in *I. scapularis*

Using BLAST, we found several groups of proteins associated with human EV biogenesis including Rab-GTPases, SNARE, ESCRT-dependent and -independent pathway, and tetraspanin proteins (Figure 1, Table S2). Prior research in *I. scapularis* involved *Vamp33*, a SNARE protein important for exosome release [13], which is located at the terminus of EV biogenesis. Thus, we became interested in proteins upstream of *Vamp33*. Among the Rab-GTPases identified, we found only one sequence with identity to both mammalian *rab27a* and *rab27b* in the *I. scapularis* genome (Figure 1, Table S2). Given this observation, we chose to examine the tick *rab27* gene.

**Fig. 1.**
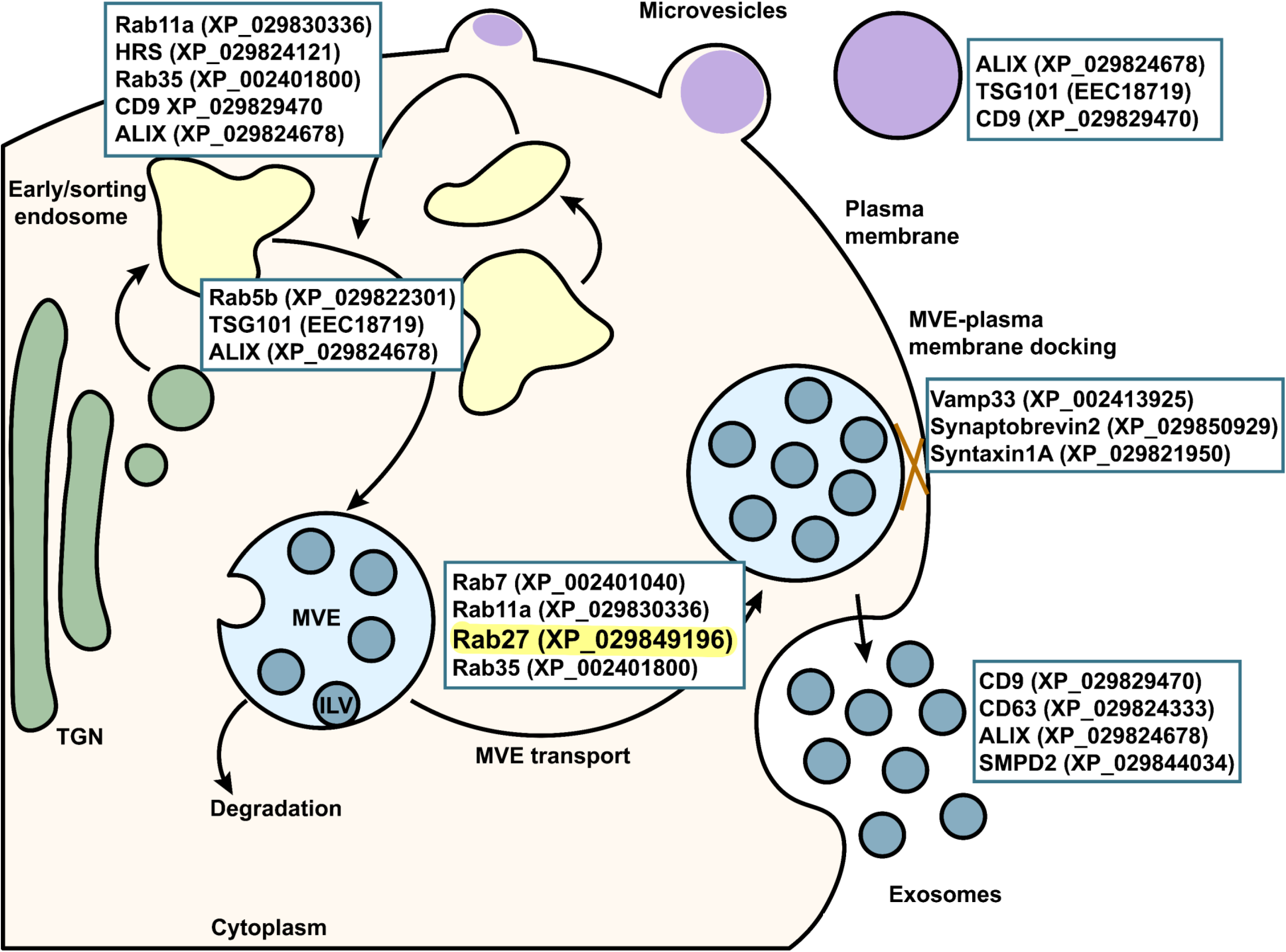
*In-silico* analysis of EV biogenesis in the *I. scapularis* genome. Sequences from proteins involved in mammalian EV biogenesis were input into the NCBI Basic Local Alignment Search Tool for proteins (BLASTp) against the *I. scapularis* genome and arranged based on mammalian literature to construct an *in-silico* pathway in ticks. Accession numbers are listed in parentheses. Rab27 is highlighted.

### Importance of Rab27 for EV biogenesis in *I. scapularis*

Previous work has shown that the tick-borne pathogens, *A. phagocytophilum*, *A. marginale*, and *Ehrlichia chaffeensis* associate with several Rab GTPases [23, 24, 25, 26, 27]. Furthermore, recent findings show that Rab27 is important for the egress of *A. phagocytophilum* from mammalian cells [28]. In mammals, Rab27a and Rab27b participate in different processes of the EV biogenesis pathway. Rab27a aids in moving the MVE to the plasma membrane for docking, and Rab27b is involved with docking [16]. Nevertheless, what role tick Rab27 plays during tick feeding or pathogen acquisition is unclear.

We opted to investigate the function of Rab27 during the formation of EVs in the tick. Small interfering RNA (siRNA) is a widely used method to knock down tick genes. Thus, we nucleofected tick ISE6 cells with tick Rab27 siRNA or scrambled RNA (scRNA) (Figure 2A). After incubating the cells for 72 hours, we removed the standard medium and replaced it with a vesicle free medium. At 24 hours, we collected vesicles as described above. We then measured their size and concentration with Nano-particle tracking analysis (NTA). We observed that the total release of vesicles in tick cells did not change between silenced and scrambled treatments (Figure 2B). However, the size range of the Rab27 silenced vesicles shifted to larger vesicles compared to the scrambled treatment (Figure 2C). The sc*-rab27*-treated tick cells mostly produced vesicles between 0-300 nm. Conversely, si-*rab27*-treated tick cells produced vesicles up to 600 nm in size and less in the 0-300 nm range. Collectively, the shift in size indicated that Rab27 is involved in tick EV biogenesis.

**Fig. 2.**
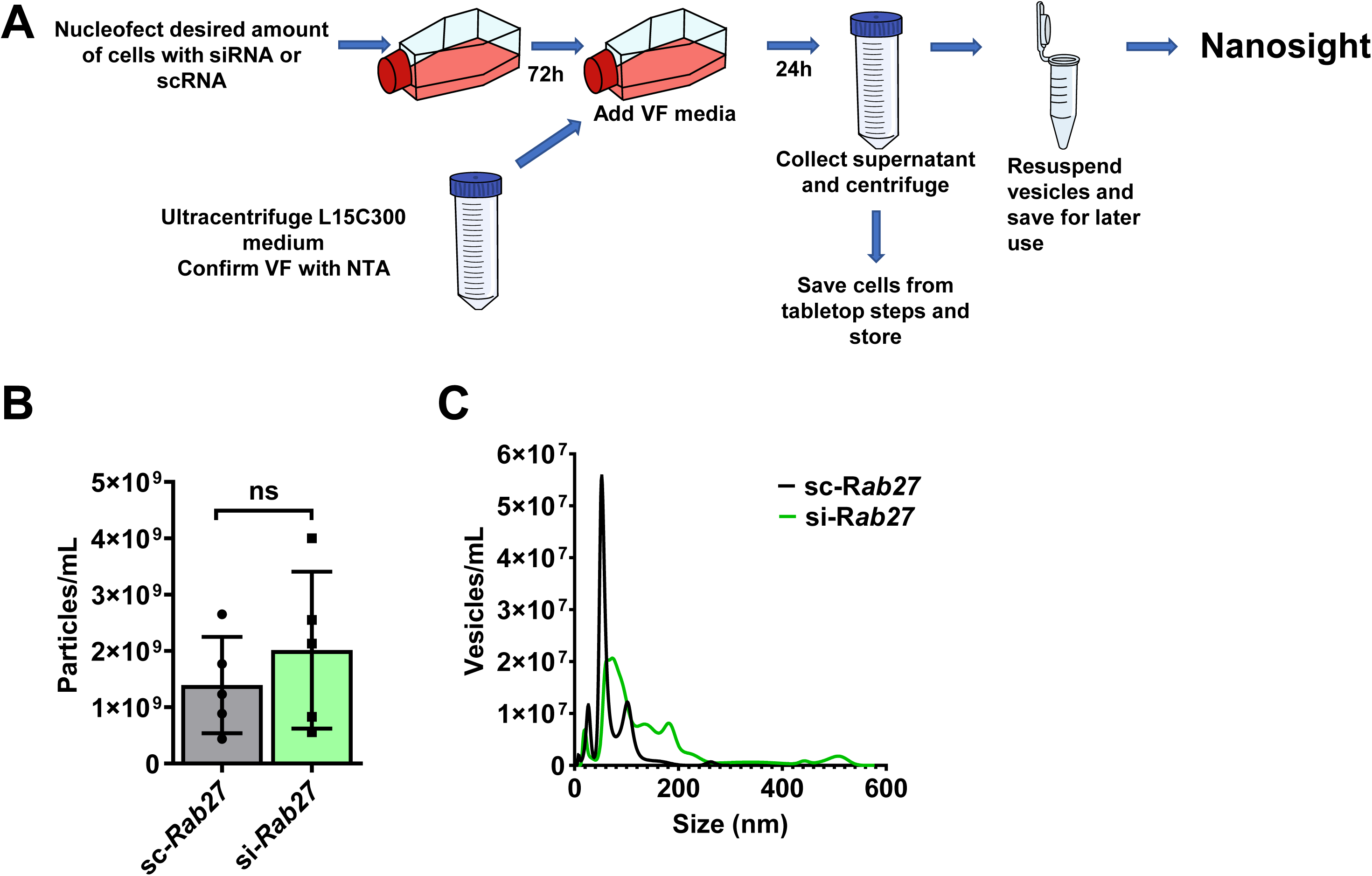
*rab27* silencing shifts the size of EVs being released by tick cells. **A** Experimental schematic for vesicle collection from tick cells. Tick ISE6 cells were nucleofected with small interfering (si-*rab27*)(green) or scrambled (sc-*rab27*)(gray) RNA for tick *rab27*. After 72-hours, the cell culture media was replaced with vesicle free (VF) media. 24-hours later, vesicles were collected through a series of tabletop and ultracentrifuge steps and measured through the nanosight machine. **B** The total vesicle amount released by tick cells does not change between the si-*rab27* or sc-*rab27* treatments. Statistical significance was evaluated by an unpaired, two-tailed *t* test. Mean ± standard deviation (SD) is plotted. ns not significant, *p*>0.05. **C** Distribution of vesicles collected from sc-*rab27* treated cells and compared to the si-*rab27* treatment ranging from 0 nm to 600 nm.

### Silencing *Rab27* impacts *I. scapularis* feeding

Next, we wanted to determine the effect of *rab27* silencing *in vivo*. We microinjected *I. scapularis* nymphs with either si-*rab27* or sc-*rab27* (Figure 3A). After 24 hours, ticks were placed on C57BL/6 mice for three days. Ticks that did not attach and fell from the mice were recovered from the water trap and counted each day. On the third day, we removed ticks that remained attached to the mouse, weighed them, and performed RNA extraction then qPCR to confirm silencing. We observed significant silencing in the RNAi treated ticks when compared to the scrambled controls (Fig 3B). Importantly, although *rab27* silencing did not affect tick attachment (Figure 3C), *rab27* silenced ticks weighed less than the scrambled counterparts (Figure 3D), indicating that Rab27 is important for *I. scapularis* hematophagy.

**Fig. 3.**
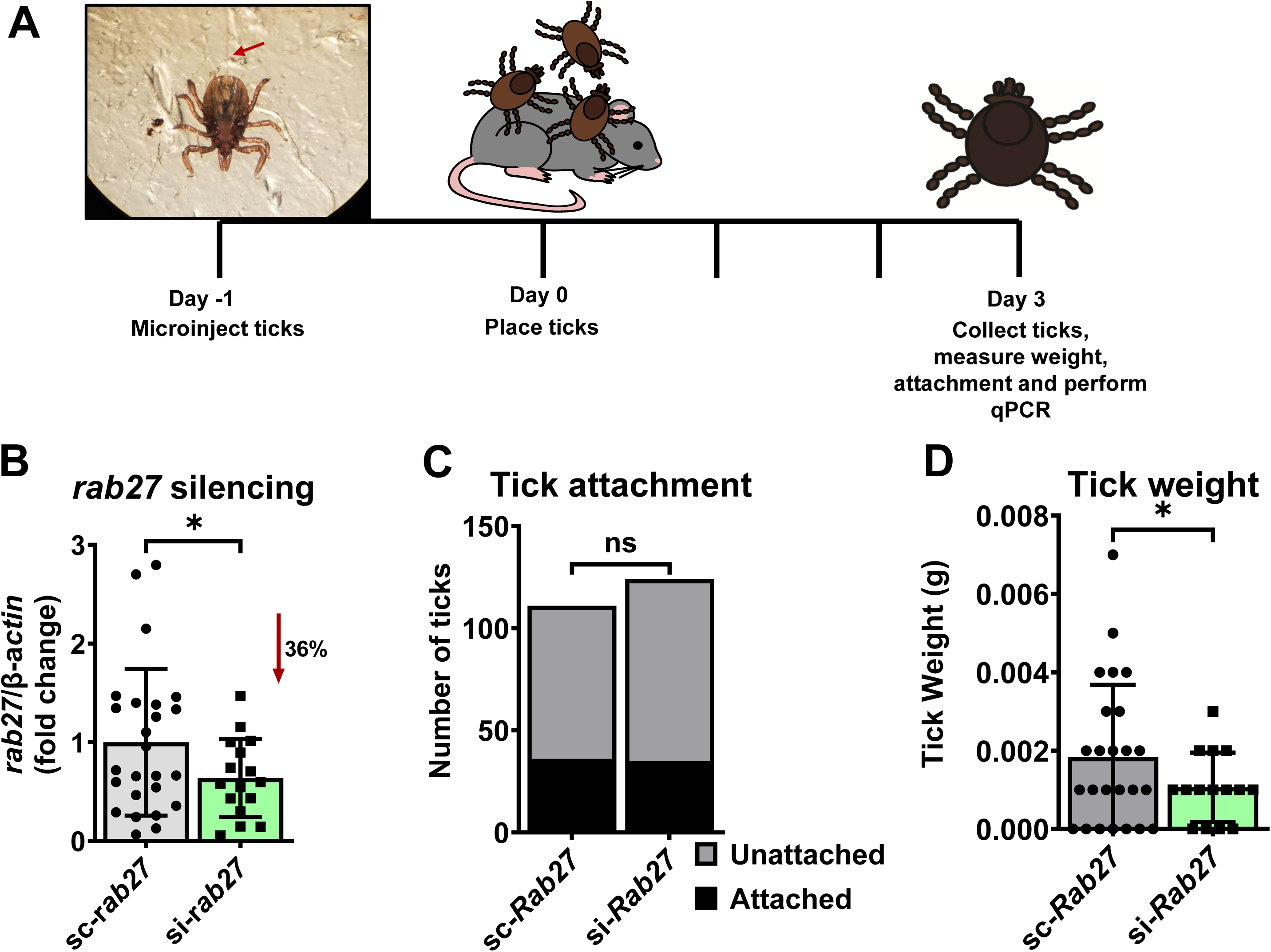
*rab27* silencing affects the fitness of *I. scapularis* nymphs during hematophagy. **A** *I. scapularis* nymphs were microinjected with small interfering (si-*rab27*) or scrambled (sc-*rab27*) RNA, incubated for 24 hours, then placed on uninfected mice. **B** *rab27* expression relative to actin after silencing. Mean ± standard deviation (SD) is plotted. Statistical significance was evaluated by an unpaired, two-tailed *t* test with Welch’s correction. \**p*<0.05 **C** *rab27* silencing effect on tick attachment. A Fisher’s exact test was performed to determine statistical differences. ns not significant, *p*>0.05. **D** Tick weight after *rab27* silencing. Mean ± SD is plotted. Statistical significance was evaluated by an unpaired, two-tailed *t* test with Welch’s correction. \**p*<0.05.

### rab27 silencing reduces acquisition of A. phagocytophilum by I. scapularis

Intracellular bacteria co-opts Rab GTPases to create a niche for their survival. Prior research has also shown that ISE6 cells infected with *A. phagocytophilum* produce an increased number of vesicles compared to uninfected cells [13]. Given that Rab27 is important for tick vesicle production, feeding, and is involved in the lifecycle of some intracellular bacteria, we explored how silencing of *rab27* affected pathogen acquisition. C57BL/6 mice were intraperitoneally injected with 10^7^ *A. phagocytophilum* seven days before tick infestation (Figure 4A). One day prior to tick placement, *I. scapularis* nymphs were microinjected with either si-*rab27* or sc-*rab27*. After a 24-hour incubation, we placed the microinjected ticks on the infected mice and allowed them to feed for three days. Unattached ticks were recovered from the water trap daily. After three days, the attached ticks were removed, weighed, and used for RNA extraction and qPCR determination of silencing. qPCR analysis detected significant levels of silencing when compared to the scrambled controls (Figure 4B). However, unlike the ticks fed on uninfected mice, significantly less *rab27* silenced ticks remained attached to the infected mice compared to the *rab27* scrambled treatment (Figure 4C). Like our previous results, *rab27* silencing led to decreased blood meal intake (Figure 4D), which resulted in a significant reduction on *A. phagocytophilum* relative levels than the scrambled ticks (Figure 4E). These results show that when *I. scapularis* feeds on an infected host, Rab27 is important for tick fitness, including attachment. It remains unclear whether the decrease observed in *A. phagocytophilum* burden was due to the diminished hematophagy in Rab27-silenced ticks or a direct effect on pathogen establishment.

**Fig. 4.**
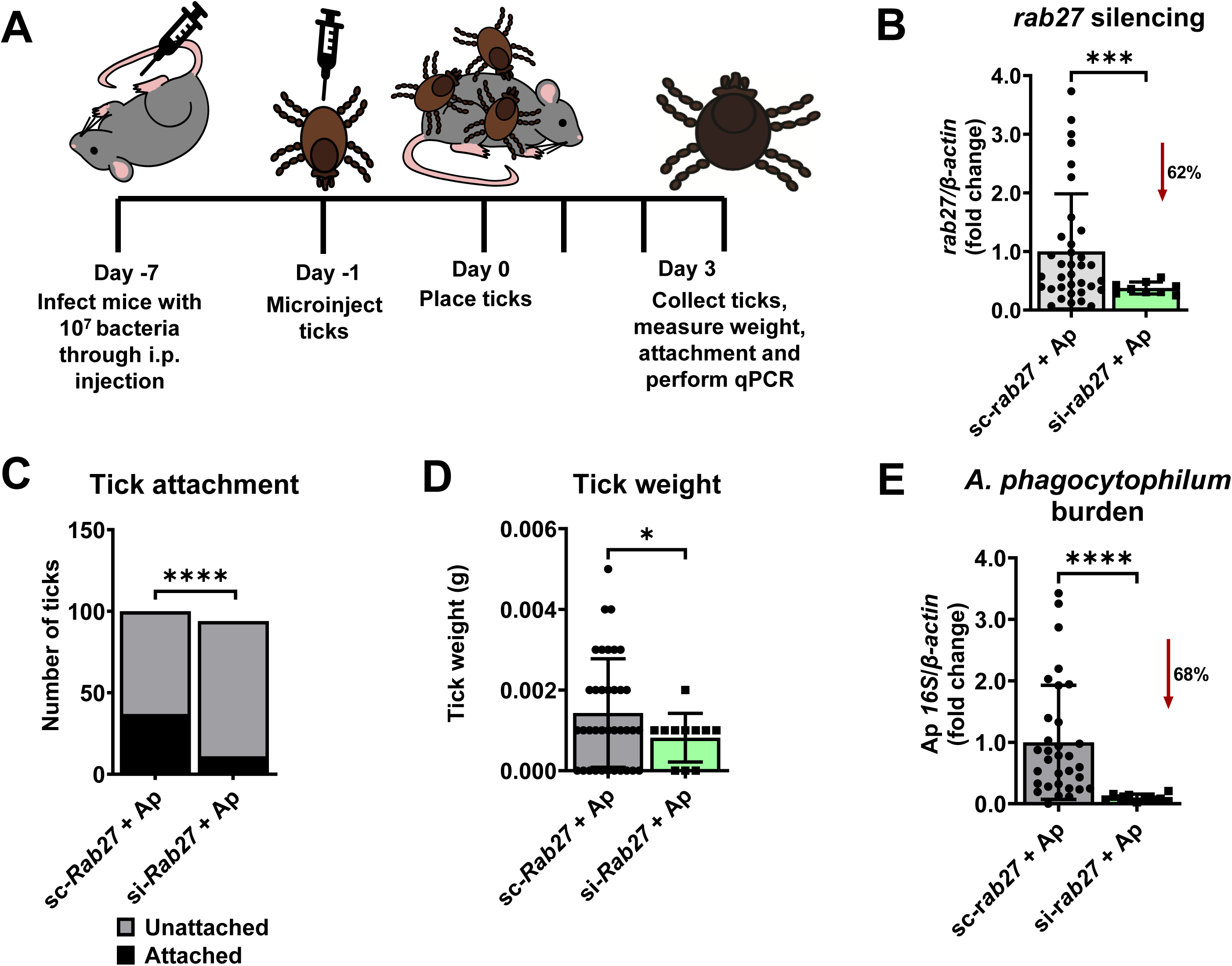
*rab27* silencing impacts *A. phagocytophilum* acquisition during blood feeding of *I. scapularis* nymphs. **A** C57BL/6 mice were intraperitoneally (i.p.) injected with 10^7^ *A. phagocytophilum* (Ap) 7 days prior to tick placement. Ticks were microinjected with small interfering (si-*rab27*) or scrambled (sc-*rab27*) RNA, incubated for 24 hours, then placed on infected mice. Ticks were recovered daily from a water trap. Remaining ticks attached to the mouse at day 3 were removed by forceps. **B** *rab27* silencing in *I. scapularis* ticks placed on Ap infected mice. Mean ± standard deviation (SD) is plotted. Statistical significance was evaluated by an unpaired, two-tailed *t* test with Welch’s correction. \*\*\**p*<0.0005 **C** Impact of *rab27* silencing on tick attachment. Fisher’s exact test was performed to determine statistical differences. \*\*\*\**p*<0.0001. **D** Effect of *rab27* silencing on tick weight. Statistical significance was evaluated by an unpaired, two-tailed *t* test with Welch’s correction. \**p*<0.05 **E** Impact of *rab27* silencing on Ap acquisition. Mean ± SD is plotted. Statistical significance was evaluated by an unpaired, two-tailed *t* test with Welch’s correction. \*\*\*\**p*<0.0001.

## Discussion

Ticks are responsible for most vector-borne disease in the United States. Due to their multiple life stages, feeding requirements, and limited genetic tools, uncovering their cellular biology has been a challenge. After silencing *rab27* in ISE6 cells, we observed a shift in vesicle size showing that a protein known to be involved in EV biogenesis in mammals is important for *I. scapularis* EV biogenesis and fitness. Mammals have two isoforms of Rab27 whereas *I. scapularis* ticks only have one present in their genome. In mammals, Rab27a and Rab27b have roles in separate stages of EV biogenesis [18]. Rab27a orchestrates the movement of the MVE to the plasma membrane of the cell. Rab27b is important for the docking of the MVE to the plasma membrane prior to exosome release [18]. Despite the change in vesicle size, it remains unclear if the tick Rab27 protein functions in a similar manner to its roles identified in mammals. It is also conceivable that the tick Rab27 protein may function distinctly from what has been described in mammalian literature. Further research will need to be completed to determine tick Rab27’s specific role in EV biogenesis.

We observed that Rab27 is important for proper tick feeding and acquisition of *A. phagocytophilum*. Because silencing *rab27* results in an alteration of vesicle size, it is plausible that the vesicles do not perform their function during tick feeding correctly resulting in reduced fitness. Alternatively, the cargo in the vesicle after *rab27* silencing may have changed, impacting vesicle function and tick fitness. We have not distinguished if the reduced acquisition of *A. phagocytophilum* was due to the impediment of tick feeding after *rab27* silencing or, alternatively, if Rab27 is needed for the life cycle of *A. phagocytophilum* in ticks. Rab27 is needed for the egress of *A. phagocytophilum* in mammalian cells [28]. Additionally, other tick-borne microbes such as *E. chaffeensis* utilize Rabs during infection by recruiting them to the occupied vacuole and delaying endosome maturation [27, 29]. Whether this manipulation occurs in ticks remains elusive Furthermore, we observed that that tick attachment was not affected when silencing *rab27* and feeding on uninfected mice. However, when *rab27* silenced ticks were fed on *A. phagocytophilum* infected mice, silenced ticks attached significantly less than scrambled treated ticks. Prior research from our lab showed that when tick cells were infected with *A. phagocytophilum*, they secreted an increased EVs compared to uninfected or *B. burgdorferi* treated cells [13]. Since silencing *rab27* disrupts EV biogenesis and *A. phagocytophilum* causes an increase of EV production, the effects of *rab27* silencing may become more pronounced when feeding ticks on infected mice, thus resulting in an attachment difference. The molecular interactions between Rab27 and *A. phagocytophilum* were not defined in the tick *I. scapularis*. However, future research could uncover an important relationship between tick EV biogenesis proteins and tick-borne diseases.

With recent advancements in the technology available to the vector-borne disease community, such as Clustered Regularly Interspaced Palindromic Repeats (CRISPR) in ticks [30], and ectopic expression in tick cells [31], future work can aim to unravel Rab27’s exact mechanism of action as well as its interaction with important tick-borne pathogens, such as *A. phagocytophilum* [32]. Furthermore, it is unclear whether Rab27 is important for tick fitness in other species. Research regarding vesicle formation in ticks and their importance for acquisition or transmission of tick-borne disease could reveal future targets to mitigate the spread of illnesses unraveling a previous unexplored aspect of vector biology. Overall, we have shown that Rab27 is important for tick EV biogenesis and fitness in *I. scapularis.* We shed light on a previously unstudied aspect of tick biology.

## Conclusions

Disrupting *rab27* impairs tick feeding and *A. phagocytophilum* acquisition during a blood meal. Additionally, silencing *rab27* in tick cells results in a shift of extracellular vesicle size. Rab27 appears to play a role in tick EV biogenesis. Future research and technological advancements will allow for further study into tick biology, the specific role of Rab27, and its importance for tick-borne disease transmission.

## Supplementary Information

**Additional file 1:** Table S1. The primers used in this study.

**Additional file 2:** Table S2. Alignment of protein sequences related to EV biogenesis.

**Additional file 3:** Table S3. Reagents used in this study.

## Abbreviations

EV: extracellular vesicle
ESCRT: endosomal sorting complex required for transport
SNARE: soluble N-ethylmaleimide-sensitive factor attachment receptor
ILVs: intraluminal vesicles
MVE: multivesicular endosome
BLASTp: Basic Local Alignment Search Tool for proteins
NTA: nanoparticle tracking analysis
siRNA: small interfering RNA
scRNA: scrambled RNA
i.p.: intraperitoneally
d.p.i.: days post infection;

## Declarations

Not applicable.

## Acknowledgements

The authors acknowledge the Pedra laboratory for providing scientific guidance and advice. We also thank Ulrike G. Munderloh (University of Minnesota) for supplying the ISE6 cells line.

## Availability of data and material

All datasets have been included with this article.

## Authors’ contributions

This project was developed by LRB and supervised by JHFP. LRB performed the research and wrote the paper with contributions from NS, LM, LMV, AJO, FECP, DKS, ASOC, and JHFP.

## Ethics approval and consent to participate

All mouse experiments were performed according to the protocols approved by the Animal Care and Use Committee (IACUC number 01121014) at the University of Maryland School of Medicine and complied with NIH guidelines (Office of Laboratory Animal Welfare [OLAW] assurance number A3200-01).

## Consent for publication

Not applicable.

