## Supplementary Tables for "Rab27 in tick extracellular vesicle biogenesis and infection"

Table S1. Primers used in this study.

| *Target* | *Type* | *Start Position (mRNA)* | *siRNA/ Primer* | *Strand* | *Primer Sequence* | *Accession Number* |
|---|---|---|---|---|---|---|
| *I. scapularis rab27* | siRNA | 325 | *Is_rab27* Si F_325 | Forward | GCTTCCTCTACCAGTACAC | XM_042289797.1 |
| | | | *Is_rab27* Si R_325 | Reverse | GTGTACTGGTAGAGGAAGC | |
| | scrambled | | *Is_rab27* Sc F_325 | Forward | GCCCTCACGTTCCATACAC | |
| | | | *Is_rab27* Sc R_325 | Reverse | TGTATGGAACGTGAGGGC | |
| | siRNA | 325 | *Is_rab27* Si F_325 | Forward | AAGCTTCCTCTACCAGTACACCCTGTCTC | |
| | | | *Is_rab27* Si R_325 | Reverse | AAGTGTACTGGTAGAGGAAGCCCTGTCTC | |
| | scrambled | | *Is_rab27* Sc F_325 | Forward | AAGCCCTCACGTTCCATACACCTGTCTC | |
| | | | *Is_rab27* Sc R_325 | Reverse | AATGTATGGAACGTGAGGGCCCTGTCTC | |
| | qRT-PCR | 382 | *Is_rab27*qR_F | Forward | GCATAGACTTCAGGGAGAAGAG | |
| | | 918 | *Is_rab27*qR_R | Reverse | CAACTGCATCCGTTATTGGG | |
| *I. scapularis actin* | qRT-PCR | 896 | *Is_actin* P2-F* | Forward | GGTCATCACAATCGGCAAC | XM_029977298.4 |
| | | 1003 | *Is_actin* P2-R* | Reverse | ATGGAGTTGTACGTGGTCTC | |
| *A. phagocytophilum 16S* | qRT-PCR | 116 | *Anaplasma 16S* F* | Forward | GGTGAGTAATGCATAGGAATC | NC_007797.1 |
| | | 223 | *Anaplasma 16S* R* | Reverse | GCTCATCTAATAGCGATAAATC | |

*Oliva Chavez, *et al.* 2021; DOI: 10.1038/s41467-021-23900-8

Table S2. Proteins associated with human EV biogenesis compared to *Mus musculus*, *Drosophila melanogaster* and *I. scapularis*

| | *Species* | *Target* | *Coverage* | *E-value* | *Identity* | *Accession Number* |
|---|---|---|---|---|---|---|
| *Rabs* | *Homo sapiens* | Rab5 | | | | AAB08927 |
| | *Mus musculus* | ras-related protein Rab-5C isoform 1 | 100% | 5.00E-160 | 98.15% | NP_077776 |
| | *Drosophila melanogaster* | Rab5, isoform G | 98% | 2.00E-112 | 74.65% | NP_001259925 |
| | *Ixodes scapularis* | ras-related protein Rab-5B | 100% | 5.00E-123 | 81.45% | XP_029822301 |
| | *Homo sapiens* | Rab7 | | | | AAD02565 |
| | *Mus musculus* | ras-related protein Rab7a | 100% | 2.00E-153 | 99.03% | NP_001280581 |
| | *Drosophila melanogaster* | Rab7, isoform B | 100% | 2.00E-119 | 75.85% | NP_001247276 |
| | *Ixodes scapularis* | ras-related protein Rab7 | 97% | 1.00E-126 | 83.17% | XP_002401040 |
| | *Homo sapiens* | Rab11a | | | | CAG38732 |
| | *Mus musculus* | ras-related protein Rab11-A | 100% | 2.00E-163 | 100.00% | NP_059078 |
| | *Drosophila melanogaster* | Rab11, isoform B | 99% | 5.00E-134 | 85.51% | NP_477170 |
| | *Ixodes scapularis* | ras-related protein Rab11-A | 99% | 3.00E-130 | 82.87% | XP_029830336 |
| | *Homo sapiens* | Rab27a | | | | CAG38735 |
| | *Mus musculus* | ras-related protein Rab-27A | 100% | 2.00E-160 | 95.93% | NP_001288159 |
| | *Drosophila melanogaster* | Rab27, isoform C | 82% | 2.00E-85 | 63.74% | NP_569921 |
| | *Ixodes scapularis* | ras-related protein Rab-27A | 86% | 1.00E-98 | 69.93% | XP_040067262.1 |
| | *Homo sapiens* | Rab27b | | | | AAM21102 |
| | *Mus musculus* | ras-related protein Rab-27B isoform 1 | 100% | 4.00E-157 | 94.50% | NP_001076022 |

| | | | | | | |
|---|---|---|---|---|---|---|
| | *Drosophila melanogaster* | Rab27, isoform C | 83% | 1.00E-79 | 61.45% | NP_569921 |
| | *Ixodes scapularis* | ras-related protein Rab-27A | 87% | 5.00E-91 | 64.92% | XP_040067262.1 |
| | *Homo sapiens* | Rab35 | | | | CAG38725 |
| | *Mus musculus* | ras-related protein Rab-35 | 100% | 1.00E-149 | 100.00% | NP_937806 |
| | *Drosophila melanogaster* | Rab35, isoform B | 100% | 2.00E-100 | 71.92% | NP_001188678 |
| | *Ixodes scapularis* | ras-related protein Rab-35 | 100% | 2.00E-115 | 77.61% | XP_002401800 |
| | *Homo sapiens* | Syntaxin 1A | | | | AAA53519 |
| *SNAREs* | *Mus musculus* | syntaxin-1A isoform 1 | 100% | 0 | 98.26% | NP_058081 |
| | *Drosophila melanogaster* | syntaxin 1A, isoform C | 99% | 3.00E-139 | 70.49% | NP_001303545 |
| | *Ixodes scapularis* | syntaxin-1A | 91% | 2.00E-130 | 71.97% | XP_029821950 |
| | *Homo sapiens* | Synaptobrevin 2 | | | | AAF15551 |
| | *Mus musculus* | vesicle associated membrane protein 2 | 100% | 5.00E-81 | 99.14% | AAB62931 |
| | *Drosophila melanogaster* | neuronal synaptobrevin, isoform K | 89% | 6.00E-46 | 66.36% | NP_001261270 |
| | *Ixodes scapularis* | vesicle-associated membrane protein 3 isoform X1 | 65% | 3.00E-36 | 76.32% | XP_029850929 |
| | *Homo sapiens* | 33kDa Vamp-associated protein | | | | AAF72105 |
| | *Mus musculus* | vesicle-associated membrane protein-associated protein A isoform 2 | 100% | 8.00E-176 | 96.69% | NP_038961 |
| | *Drosophila melanogaster* | VAMP-associated protein 33kDa, isoform A | 78% | 5.00E-42 | 44.04% | NP_996348 |
| | *Ixodes scapularis* | vesicle-associated membrane protein-associated protein B isoform X1 | 98% | 3.00E-70 | 44.87% | XP_002413925 |
| *ESCRT-dependent* | *Homo sapiens* | Hrs (ESCRT-0) | | | | BAA23366 |

| | | | | | | |
|---|---|---|---|---|---|---|
| | *Mus musculus* | hepatocyte growth factor-regulated tyrosine kinase substrate isoform 1 | 100% | 0.00E+00 | 93.44% | NP_001152800 |
| | *Drosophila melanogaster* | hepatocyte growth factor regulated tyrosine kinase substrate, isoform B | 72% | 6.00E-91 | 48.09% | NP_525099 |
| | *Ixodes scapularis* | hepatocyte growth factor-regulated tyrosine kinase substrate | 72% | 0.00E+00 | 52.83% | XP_029824121 |
| | *Homo sapiens* | TSG101 (ESCRT-1 subunit TSG101) | | | | Q99816 |
| | *Mus musculus* | tumor susceptibility gene 101 protein isoform 1 | 10% | 0.00E+00 | 94.63% | NP_068684 |
| | *Drosophila melanogaster* | tumor susceptibility gene 101 | 98% | 7.00E-134 | 50.61% | NP_524120 |
| | *Ixodes scapularis* | tumor susceptibility protein, putative, partial | 95% | 1.00E-132 | 49.74% | EEC18719 |
| | *Homo sapiens* | ALIX; programmed cell death 6 interacting protein isoform 2 | | | | NP_001155901 |
| | *Mus musculus* | programmed cell death 6-interacting protein isoform 1 | 100% | 0.00E+00 | 94.39% | NP_001158149 |
| | *Drosophila melanogaster* | ALG-2 interacting protein X | 100% | 0.00E+00 | 41.38% | NP_651582 |
| | *Ixodes scapularis* | programmed cell death 6-interacting protein | 94% | 0.00E+00 | 44.95% | XP_029824678 |
| *ESCRT-independent* | *Homo sapiens* | sphingomyelin phosphodiesterase 2, neutral membrane (SMPD2) | | | | EAW48346 |
| | *Mus musculus* | sphingomyelin phosphodiesterase 2 | 99% | 0.00E+00 | 77.14% | NP_033239 |
| | *Drosophila melanogaster* | neutral sphingomyelinase | 65% | 5.00E-50 | 34.98% | NP_647790 |
| | *Ixodes scapularis* | sphingomyelin phosphodiesterase 2 isoform X1 | 65% | 1.00E-72 | 40.62% | XP_029844034 |
| *Tetraspanins* | *Homo sapiens* | CD9 antigen isoform 1 | | | | NP_001760 |
| | *Mus musculus* | CD9 antigen | 100% | 2.00E-150 | 89.04% | NP_031683 |
| | *Drosophila melanogaster* | tetraspanin 96F, isoform C | 98% | 2.00E-22 | 23.05% | NP_001262977 |

| | | | | | | |
|---|---|---|---|---|---|---|
| | *Ixodes scapularis* | CD9 antigen isoform X2 | 98% | 3.00E-38 | 32.84% | XP_029829470 |
| | *Homo sapiens* | CD63 antigen isoform A | | | | NP_001771 |
| | *Mus musculus* | CD63 antigen | 100% | 2.00E-139 | 79.41% | NP_001036045 |
| | *Drosophila melanogaster* | tetraspanin 39D | 97% | 5.00E-32 | 28.33% | NP_523612 |
| | *Ixodes scapularis* | 23 kDa integral membrane protein | 100% | 2.00E-38 | 32.50% | XP_029824333 |
| | *Homo sapiens* | CD81 antigen isoform 2 | | | | NP_001284578 |
| | *Mus musculus* | CD 81 antigen, isoform CRA_a | 100% | 2.00E-104 | 89.70% | EDL18193 |
| | *Drosophila melanogaster* | tetraspanin 96F, isoform C | 95% | 6.00E-13 | 25.26% | NP_001262977 |
| | *Ixodes scapularis* | CD9 antigen isoform X2 | 97% | 6.00E-14 | 26.60% | XP_029829470 |

Table S3. Reagents used in this study.

| *Cell Medium* | **Source** | **Identifier** | **Dilution/Concentration** |
|---|---|---|---|
| Leibovitz's L-15 Medium, powder | Gibco | 41300039 | N/A |
| L-aspartic acid | Millipore-Sigma | 11189 | 0.449 g/L |
| L-glutamine | Millipore-Sigma | G8540 | 0.500 g/L |
| L-proline | Millipore-Sigma | 81709 | 0.450 g/L |
| L-glutamic acid | Millipore-Sigma | 49449 | 0.250 g/L |
| alpha-ketoglutaric acid | Millipore-Sigma | K1128 | 0.449 g/L |
| Sodium hydroxide | Millipore-Sigma | S8045 | 10 N |
| D-glucose | Millipore-Sigma | G7021 | 18.018 g/L |
| FBS (USDA approved; for tick media) | Millipore-Sigma | F0926-500ML | 0.1 |
| Bacto™ Tryptose Phophate Broth | BD | 260300 | 5% |
| Lipoprotein Concentrate | MP Biomedicals | 191476 | 0.10% |
| RPMI-1640 Medium With L-Glutamine | Quality Biological | 112-025-101 | N/A |
| Fetal Bovine Serum (for HL-60s media) | Gemini Bio-Products | 100-106 | 0.1 |
| GlutaMax | Gibco | 35050-061 | 0.01 |
| *Materials* | | | |
| Cellstar® cell culture flasks, 25 cm2 Greiner bio-one 690-160 N/A | Greiner bio-one | 690-160 | N/A |
| T-25 Vented Flasks | CytoOne | CC7682-4825 | N/A |

| | | | |
|---|---|---|---|
| 1.5 ml microcentrifuge tubes | Thermo Scientific | 1148T71 | N/A |
| 15 ml conical screw cap tubes | USA Scientific | 5618-8261 | N/A |
| 50 ml conical screw cap tubes | USA Scientific | 5622-7270 | N/A |
| 500 ml vacuum filter/storage bottle system, 0.2 µm | Corning | 430773 | N/A |
| Rnase-free disposable pellet pestles | Thermo Scientific | 12-141-368 | N/A |
| Fisherbrand™ Superfrost™ Plus Microscope Slides | Thermo Scientific | 12-550-15 | N/A |
| White Filter Paper for CytoSep™ Single Funnel | Simport Scientific | M965FW | N/A |
| *Reagents* | | | |
| 2X Universal SYBR Green Fast qPCR Mix | Abclonal | RK21203 | 1X |
| Ethyl alconol, Pure; 200 proof for molecular biology | Millipore-Sigma | E7023-1L | 1:01 |
| TRIzol reagent | Ambion | 15596018 | N/A |
| 10X Phosphate-buffered saline (PBS) | Quality Biological | 119-069-131 | 1X |
| Chloroform | Millipore-Sigma | 288306 | 1 |
| HyClone™ Water, Molecular Biology Grade | Cytiva | SH3053801 | N/A |
| Commercial assays | | | |

| | | | |
|---|---|---|---|
| Pure Link RNA mini kit | Ambion | 12183025 | N/A |
| Silencer™ siRNA Construction Kit | Thermo Scientific | AM1620 | N/A |
| Verso cDNA Synthesis Kit | Thermo Scientific | AB-1453B | N/A |
| *Mycoplasma* testing kit | Southern Biotech | 13100-01 | N/A |
| SF Cell Line 4D-NucleofectorTM X Kit L | Lonza Bioscience | V4XC-2012 | N/A |
| Richard-Allan Scientific™ Three-Step Stain Set | Thermo Scientific | 3300 | N/A |
| *Organisms* | | | |
| *Ixodes scapularis* nymph ticks | Jonathan Oliver and Ulrike Munderloh, University of Minnesota | N/A | N/A |
| *Ixodes scapularis* nymph ticks | Tick Lab, Oklahoma State University | N/A | N/A |
| C57BL6J (WT) mice | University of Maryland | N/A | N/A |
| *A. phagocytophilum* HZ | Ulrike Munderloh, University of Minnesota | N/A | N/A |
| Cell lines | | | |
| *I. scapularis* ISE6 cells | Ulrike Munderloh, University of Minnesota | ISE6 | N/A |
| HL-60 | ATCC | CCL-240 | N/A |
| Equipment | | | |
| CFX96 Touch Real-Time PCR Detection System | Biorad | N/A | N/A |

| | | | |
|---|---|---|---|
| 4D-NucleofectorTM System | Lonza Bioscience | AAF-1002 | N/A |
| Epredia™ Cytospin™ 4 Cytocentrifuge | Thermo Scientific | A78300003 | N/A |
| Carl Zeiss™ Primo Star™ HAL/LED Microscope | Carl Zeiss | 415500-0057-000 | N/A |
| Nanoject III | Drummond Scientific Company | 3-000-207 | N/A |
| NanoSight NS300 | Malvern Panalytical | N/A | N/A |
| Percival I30BLL incubator | Percival | I30BLL | N/A |
| *Software* | | | |
| GraphPad Prism v10.0.3 | https://www.graphpad.com/ | N/A | N/A |
| Graphpad Quick Cals Outlier Calculator | https://www.graphpad.com/quickcalcs/Grubbs1.cfm | N/A | N/A |
| BLOCK-iT RNAi designer | https://rnaidesigner.thermofisher.com/rnaiexpress/ | N/A | N/A |
| InvivoGen siRNA Wizard Software | https://www.invivogen.com/sirnawizard/scrambled.php | N/A | N/A |
| NIH Basic Local Alignment Search Tool for proteins | https://blast.ncbi.nlm.nih.gov/Blast.cgi?PAGE=Proteins | N/A | N/A |
| NIH Basic Local Alignment Search Tool for nucleotides | https://blast.ncbi.nlm.nih.gov/Blast.cgi?PROGRAM=blastn&PAGE_TYPE=BlastSearch&BLAST_SPEC=&LINK_LOC=blasttab&LAST_PAGE=blastp | N/A | N/A |
| PrimerQuest Tool | https://www.idtdna.com/PrimerQuest/Home/Index | N/A | N/A |
